# Altered Reward Processing in Abstinent Dependent Cannabis Users: Social Context Matters

**DOI:** 10.1101/278044

**Authors:** Kaeli Zimmermann, Keith M. Kendrick, Dirk Scheele, Wolfgang Dau, Markus Banger, Wolfgang Maier, Bernd Weber, Yina Ma, René Hurlemann, Benjamin Becker

## Abstract

Public perception of cannabis as relatively harmless, alongside claimed medical benefits, have led to moves towards its legalization. Yet, long-term consequences of cannabis dependence, and whether they differ qualitatively from other drugs, are still poorly understood. A key feature of addictive drugs is that chronic use leads to adaptations in reward processing, blunting responsivity to the substance itself and other rewarding stimuli. Against this background, the present study investigated whether cannabis dependence is associated with reductions in hedonic representations by measuring behavioral and neural responses to social reward in 23 abstinent cannabis-dependent men and 24 matched non-using controls. In an interpersonal pleasant touch fMRI paradigm, participants were led to believe they were in physical closeness of or touched (CLOSE, TOUCH) by either a male or female experimenter (MALE, FEMALE), allowing the assessment of touch- and social context-dependent (i.e. female compared to male social interaction) reward dynamics.

Upon female compared to male touch, dependent cannabis users displayed a significantly attenuated increase of reward experience compared to healthy controls. Controls responded to female as compared to male interaction with increased striatal activation whereas cannabis users displayed the opposite activation pattern, with stronger alterations being associated with a higher lifetime exposure to cannabis. Neural processing of pleasant touch in dependent cannabis users remained intact.

These findings demonstrate that cannabis dependence in men is linked to similar lasting neuroadaptations in striatal responsivity to hedonic stimuli as observed for other drugs of abuse. However, reward processing deficits seem to depend on the social context.

## INTRODUCTION

Together with claimed medical benefits, perception of cannabis as less harmful than other drugs (Anthony et al., 1994) has promoted recent moves towards legalization. With long-term regular use, however, dependence risks increase, and relapse rates are comparable to other drugs (Hall and Degenhardt, 2009). Although neuroadaptations associated with cannabis use have been examined extensively, most studies focused on recreational users, or dependent users during early abstinence, a period characterized by withdrawal (Budney et al., 2003), neural recovery (Hirvonen et al., 2012) and potential residual effects of cannabis metabolites for up to 28 days (McGilveray, 2005). Functional alterations have been reported to both normalize and persist (Sneider et al., 2008) 4 weeks following cessation of cannabis use. Whether persistent neurobiological changes related to cannabis dependence are similar to those observed following chronic exposure to other drugs thus remains a subject of debate.

Current conceptualizations of addiction propose dysregulations in reward circuits leading to lasting allostatic adaptations in hedonic processing (Volkow et al., 2012; Koob, 2015). Animal models have linked the mesolimbic system, particularly striatal nodes, to acute drug reward signaling and neuroadaptations thereof are thought to drive compulsive drug seeking (Di Chiara and Imperato, 1988). Studies in human users suggest that exaggerated striatal reactivity to drug-reward cues and concomitantly reduced sensitivity for natural (non-drug) rewards (Volkow et al., 2012) contribute to the addictive process during which drug seeking becomes the central motivational drive and promote relapse (Lubman et al., 2009). This imbalance at the core of the brain’s reward circuit thus plays an important role in the behavioral maladaptations in dependent individuals.

Previous findings on non-drug reward processing in cannabis users following short abstinence remain inconsistent (Nestor et al., 2010; Jager et al., 2013; Martz et al., 2016). Residual effects of chronic cannabis use on striatal blood flow can be observed even after 72h of abstinence (Filbey et al., 2017) and, together with the use of monetary rewards, which associate with drug-cue properties, may have contributed to the inconsistencies. Moreover, is that alterations across striatal subregions in cannabis users strongly depend on the social context, such as exposure to social information (Gilman et al., 2016).

Social factors such as peers considerably influence the addictive process and predict initiation and escalation of use, and treatment success (Nikmanesh et al., 2015). In return, drug use itself profoundly affects social behavior ranging from initially enhanced sociability to social withdrawal once a dependence has been developed (McGregor et al., 2008). Therefore, social interaction deficits are increasingly recognized as core characteristics of drug use disorders (DSM 5). In line with these observations, animal models indicate lasting social impairments and reduced social interactions following chronic drug exposure (O’Shea et al., 2006) possibly rooted in deficient striatal sensitivity for social rewards (Zernig and Pinheiro, 2015). Indeed, positive social interactions engage the striatal reward system (Izuma et al., 2008) and may represent an alternative natural reward to drug use.

Pleasant interpersonal touch is a vital instrument for conveying social reward and positive social interaction (Ellingsen et al., 2016). As a powerful natural reward, the affective experience of pleasant interpersonal touch elicits activations in the brain’s reward network (Ellingsen et al., 2016). Both the hedonic experience and associated striatal response strongly depend on the social context (Kreuder et al., 2017). Specifically, increased pleasantness and striatal activity have been observed when male subjects believe touch is applied by a female as opposed to a male experimenter (Scheele et al., 2014).

The present study addressed whether cannabis dependence is associated with lasting impairments in processing of social rewards and whether these impairments depend on the social context. A pleasant interpersonal touch fMRI paradigm (Gazzola et al., 2012; Scheele et al., 2014) was employed allowing social context-dependent reward variation by making abstinent (≥28 days) cannabis-dependent men and controls believe that pleasant touch was applied by either a female or male experimenter.

Based on the proposed significance of blunted natural reward sensitivity and social impairments in drug dependence, we expected reduced hedonic experience of pleasant touch and its contextual modulation. In accordance with recent evidence for social context-dependent striatal alterations in cannabis users (Gilman et al., 2016) we furthermore expected blunted striatal coding of reward modulation induced by opposite sex as compared to same sex interaction.

## MATERIALS and METHODS

### Participants

For selection pipeline of study sample see **SI**. To control for confounding effects of hormonal fluctuations related to menstrual cycle or contraceptives on the outcome parameters, including reward-related striatal activity (Dreher et al., 2016), and dependence symptoms such as craving (Franklin et al., 2015), the present study focused on male participants (similar approach see Zimmermann et al., 2017). 23 abstinent dependent cannabis users and 24 demographically-matched non-using controls were scheduled for the assessment that included questionnaires, cognitive tests, drug urine screen and fMRI. Inclusion criteria for all participants were: 1) Age 18-35, 2) right-handedness, 3) heterosexuality and 4) a negative urine toxicology for cannabis and other illicit drugs (Drug-Screen^®^ Pipette test, Nal van Minden, Moers, Germany, Multi 7TF for amphetamines (cut-off: 500 ng/ml), cocaine (300 ng/ml), methamphetamine (500 ng/ml), THC (50 ng/ml), MDMA (300 ng/ml), opiate (300 ng/ml), methadone (300 ng/ml)) at the day of the fMRI assessment. Cannabis users were included if they fulfilled the DSM-IV criteria for cannabis dependence during the previous 18 months and agreed to abstain from cannabis in the 28 days before the assessment. At the time of enrollment, most users were still using cannabis or were in an early phase of abstinence. Cannabinoid metabolites remain in the body for up to 4 weeks after cessation (McGilveray, 2005) and withdrawal symptoms peak in the first week after last of use (Budney et al., 2003). Therefore, a minimum abstinence of 28 days was selected to allow the assessment of lasting effects, in line with comparable MRI studies (Sneider et al., 2008). Abstinence was based on self-report and negative urine toxicology. Active cannabis users were included if they were willing to abstain for 28 days and currently abstinent users were asked to maintain abstinent for the 28 days prior to fMRI assessment. One user reported having used cannabis on one occasion 14 days before the experiment, but was included due to a negative urine toxicology. Control subjects were included if their cumulative lifetime cannabis use was below 10g. Exclusion criteria for all participants were: 1) any profound DSM-IV axis I or axis II disorder, e.g. psychotic or bipolar disorders, 2) Beck Depression Inventory score (BDI-II) ≥ 20 (maximum BDI in the final sample = 15, mean scores comparable for users and controls, p > .05), 3) medical disorder, 4) current/regular medication intake, and 5) MRI-contraindications. Attention, attitude toward interpersonal touch, social interaction anxiety, anxiety, mood and relationship status (y/n) were assessed as potential confounders (details **SI**). Experience with other licit and illicit drugs was documented. Given that the co-use of other illicit substances is common in cannabis users, users with > 75 lifetime occasions of other illicit drugs were excluded. Due to high co-occurrence of cannabis and tobacco use (Agrawal et al., 2012), groups were matched for the number of tobacco smokers and use patterns. As a trade-off between confounding effects of acute nicotine and nicotine craving on striatal reward processing, all smokers underwent 1.5h of supervised abstinence before the fMRI. Users were recruited in cooperation with the Department of Addiction and Psychotherapy of the LVR Clinics Bonn (Germany). Written informed consent was obtained from all participants. The study had full ethical approval by the University of Bonn and was registered as clinical trial (NCT02711371). Procedures were in accordance with the latest revision of the Declaration of Helsinki.

### Interpersonal Touch Paradigm

An interpersonal touch fMRI paradigm with context-dependent reward variation was employed (Scheele et al., 2014; adapted from Gazzola et al., 2012). Before entering the scanner participants were introduced to a male and female experimenter that were the same throughout the study. The experiment consisted of two sessions (one male, one female), each with three conditions indicated by photographs depicting the experimenter: ‘HOME’, where the experimenter stands at 2 m distance, ‘CLOSE’, where the experimenter stands at the junction of the MRI table and opening, and ‘TOUCH’, where the experimenter administers repeated soft touch using downwards strokes to the shin of both legs (20 cm on the shin, velocity: 5 cm/s). This design allowed to vary rewarding properties and to assess two natural social reward dimensions (‘TOUCH > CLOSE’ as touch-associated reward, ‘FEMALE > MALE’ as context-dependent reward). To control for differences in physical properties of touch, only the male experimenter applied the soft strokes (details see **SI**). Following each ‘CLOSE’ and ‘TOUCH’ trial subjects rated the perceived pleasantness (1 (unhappy emoticon) ‘very unpleasant’ to 20 (happy emoticon) ‘very pleasant’, see also Scheele et al., 2014; Kreuder et al., 2017; based on the SAM non-verbal assessment for affective experience (Bradley & Lang, 1994)). All participants rated attractiveness and likeability of the experimenters on a scale from 0 (not likeable at all; not attractive at all) to 10 (very likeable; very attractive) after the experiment. Cannabis craving was assessed before and after fMRI (CCS-7; Schnell et al., 2011).

### Behavioral Data Analysis

Data was analyzed in SPSS20 (SPSS Inc., Chicago, IL, USA). Demographic and questionnaire data were analyzed using independent t-tests (for non-normal distributed data corresponding non-parametric analyses were used) and results considered significant at p< .05 (two-tailed). Median and range are reported for non-normal distributed data.

Pleasantness ratings were examined by mixed analysis of variance (ANOVA) with condition (touch vs close) and experimenter (male vs female) as within-subject factors and group (users vs controls) as between-subject factor. To more specifically address the hypothesized reduced reward dynamics in cannabis users an exploratory analysis focused on the comparison of the two conditions (female touch > male touch) that showed the strongest pleasantness increase in previous studies (Gazzola et al., 2012; Scheele et al., 2014). To this end between-group differences in the mean percent pleasantness increase between these conditions ([(pleasantness rating_FemaleTouch_ –pleasantness rating_MaleTouch_)/pleasantness rating_MaleTouch_]*100) were compared using an independent t-test. Specifically, this targeted analysis allowed to address the strongest gain in reward value and therefore appears specifically sensitive to capture reduced reward dynamics. One cannabis user was excluded due to consistently rating male touch as very aversive (consistent rating_MaleTouch_ = 1) (details see SI), resulting in n = 22 cannabis users and n = 24 controls entering the final analyses.

### fMRI Data Acquisition and Analysis

Data was acquired on a Siemens 3 Tesla system using established scanning and preprocessing procedures (**SI**). The first level model included four conditions: ‘TOUCH_Female_’, ‘CLOSE_Female_’, ‘TOUCH_Male_’, and ‘CLOSE_Male_’. ‘HOME’ served as implicit baseline and motion parameters were included as additional regressors. Condition-specific regressors were convolved with the hemodynamic response function and estimated using a general linear model (GLM). In line with the pleasantness ratings, a mixed ANOVA including the within-subject factors touch *vs* close and male *vs* female, and the between-subject factor group (users *vs* controls) was performed. The ANOVA was implemented using a partitioned error- approach and first level contrasts assessing dynamic coding of touch-associated reward (‘TOUCH > CLOSE’), context-dependent reward (‘FEMALE > MALE’), and their interaction (‘FEMALE _touch>close_ > MALE _touch>close_’). Groups were compared in SPM independent t-tests. Results were thresholded using a cluster-level FWE-correction of p < .05 (in line with recent recommendations an initial cluster-defining threshold of p < .001 was applied to data resampled at 3x3x3 mm^2^, Slotnick, 2017).

Parameter estimates were extracted from significant clusters showing group differences (contrasts: ‘FEMALE > MALE’; ‘FEMALE > baseline’, ‘MALE > baseline’). Associations between use-based measures of dependence severity (cumulative lifetime amount [z-transformed]) and recovery (days since last use [z-transformed]), as well as measures of withdrawal (BDI-II, STAI and CCS-7) with behavioral and neural indices were examined using bivariate correlation (p < .05, two-tailed).

## RESULTS

### Group Characteristics

Groups were comparable in potential confounders, including alcohol/nicotine use (**Table 1**). Cannabis users reported comparable low craving before and after the experiment (scale 7-49; pre: 19.05 ±11.37; post: 18.68 ±10.72, p = .67, dependent t-test). **Table 2** shows cannabis use parameters. Examining mood scores using an ANOVA with the within-subject factor assessment time (pre- *vs* post-experiment) and the between subject factor group (users *vs* controls) did not reveal significant differences (all p > .14). Together, craving and mood data argue against confounding effects of acute cannabis withdrawal.

**Table 1.** Group characteristics and drug use parameters. ^a^Mann-Whitney-U test, *Median(Range), ** Prescription medicinal use.

| Measure | Cannabis Users<br>M (SD) | Controls<br>M (SD) | p |
| --- | --- | --- | --- |
| Age | 23.86(3.36) | 23.67(2.88) | .83 |
| Years of education | 15.00(11.00-22.00)* | 14.50(12.00-19.00)* | .92<sup>a</sup> |
| d2 concentration performance | 196.32(41.19) | 201.75(52.95) | .70 |
| STQ mean | 1.15(0.80-1.85)* | 1.20(0.55-2.40) | .86<sup>a</sup> |
| STAI state | 33.95(8.08) | 30.54(7.49) | .14 |
| STAI trait | 35.55(8.34) | 32.46(6.92) | .18 |
| SIAS | 18.00(8.00-46.00)* | 16.00(5.00-33.00)* | .32<sup>a</sup> |
| Relationship status (N) (y/n) | (13/9) | (12/12) |  |
| Age of first nicotine use | 14.62(1.93) | 15.02(4.56) | .71 |
| Years of nicotine use | N = 21<br>9.25(1.00-18.00)* | N = 22<br>7.00(2.00-17.00)* | .29<sup>a</sup> |
| Cigarettes per day | 6.50(0-20.00)* | 10.00(1-20.00)* | .24<sup>a</sup> |
| Age of first alcohol intake | 14.00(11.00-16.00)* | 14.00(8.00-16.00)* | .34<sup>a</sup> |
| Alcohol occasions per week | 2.00(0-4.00)* | 1.00(0-4.00)* | .18<sup>a</sup> |
| Alcohol units per week | 6.00(0-46.00)* | 4.90(0-18.00)* | .66<sup>a</sup> |
| Past ecstasy use | N = 13 | N = 2 |  |
| Lifetime occasions ecstasy | 14.67(1-75)* | (1-8)* | - |
| Past cocaine use | N = 10 | N = 0 |  |
| Lifetime occasions cocaine | 5.98(1-70)* | - | - |
| Past amphetamine use | N = 13 | N = 1 |  |
| Lifetime occasions amphetamine | 20(1-75)* | 6.00 | - |
| Past hallucinogen use | N = 10 | N = 0 |  |
| Lifetime amount hallucinogen | 5.50(1-50)* | - | - |
| Past opiate use | N = 3 | N = 1 |  |
| Lifetime occasions opiate | 2.00(1.73) | 30.00** | - |
| Past cannabis use | N = 22 | N = 21 |  |
| % Lifetime cannabis dependence | 100% | 0% | - |

**Table 2.** Cannabis use parameters. *Median

| **Cannabis Use Parameter** | **Mean ± SD (range) (N = 22)** |
| --- | --- |
| **Age of first cannabis use** | 15.14 ± 1.27 (13 - 17) |
| **Days since last cannabis use** | 30.00* (14 - 500) |
| **Frequency of cannabis use (days per month)** | 27.91 ± 4.68 (14 - 30) |
| **Duration of regular cannabis use (months)** | 77.05 ± 36.56 (19 - 144) |
| **Lifetime amount of cannabis in grams** | 1503.50* (62 - 5786) |

### Perceived Attractiveness and Likability

Examination using repeated-measures ANOVAs including group (users *vs* controls) as between-subject factor and experimenter (male *vs* female) as within-subject factor revealed a main effect of experimenter for both, attractiveness (F = 37.97, p < .001) and likability (F = 15.33, p < .001), however no main or interaction effects with group (all p > .12), suggesting that the female experimenter was perceived as more attractive (female: 9.01 ±1.19; male: 5.05 ±1.95) and likable (female: 8.67 ±1.39; male: 7.68 ±1.21) across groups.

### Behavioral Results

Examining the pleasantness ratings revealed a significant main effect of condition (F_(1,44)_ = 11.61, p = .001, η ^2^ = .21) and experimenter (F_(1,44)_ = 4.84, p = .033, η^2^ = .01) as well as a significant interaction between these factors (F_(1,44)_ = 32.40, p < .001, η ^2^ = .42), however no effects involving the factor group reached significance (all p > .17). Across groups TOUCH (mean ±SD: 12.63 ±2.41) was rated as significantly more pleasant than CLOSE (11.41 ±2.74), and FEMALE presence (12.18 ±2.42) was rated as significantly more pleasant than MALE presence (11.87 ±2.22) (effect sizes comparable to Scheele et al., 2014). Post-hoc tests further revealed that female touch was rated as more pleasant than all other conditions (all p < .001). Comparing increased pleasantness experience for female relative to male touch revealed a significantly lower increase in cannabis users (mean % increase ± SD: 4.49 ±6.79) relative to controls (10.79 ±12.27; t_(44)_ = 2.13, p = .04, Cohen’s d = .64) (**Figure 1**).

**Figure 1.**
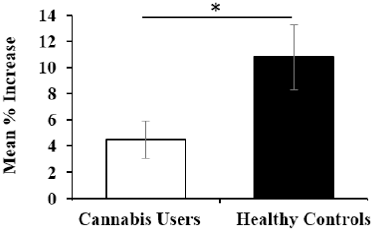
Group Differences in Mean % Increase of Pleasantness. Relative to controls, cannabis users show a significantly lower increase in pleasantness to female touch as compared to male touch. Mean % increase = [(pleasantness rating_FemaleTouch_ – pleasantness rating_MaleTouch_)/pleasantness rating_MaleTouch_]*100. Error bars indicate SEM. * p < .05.

### fMRI Results

We initially replicated previous findings (Gazzola et al., 2012; Scheele et al., 2014). The application of soft touch (‘TOUCH > CLOSE’) elicited activity in a network encompassing primary somatosensory, striatal and insula regions in controls (p < .05; see **SI Figure S1, Table S1**) possibly reflecting the sensory and rewarding properties of pleasant soft touch. Cannabis users engaged a similar network (see **Figure S1, Table S1**). The contextual modulation of pleasant touch (‘FEMALE _touch>close_ > MALE _touch>close_’) in controls revealed significant interaction effects in the right somatosensory cortex (peak at MNI 30 / -37 / 37, t_(23)_ = 5.54, k = 352, p < .001), the right posterior insula (peak at 33 / -13 / 20, t_(23)_ = 5.40, k = 72, p = .025) and the left precentral gyrus (peak at -24 / -16 / 41, t_(23)_ = 5.29, k = 223, p < .001) in accordance with previous studies (Gazzola et al., 2012; Scheele et al., 2014) and meta-analyses (Morrison, 2016) on the involvement of these regions in affective modulation of touch. For cannabis users no significant interaction effects were observed.

Groups did not differ significantly in touch-related processing (‘TOUCH > CLOSE’) and its contextual modulation (‘FEMALE _touch>close_ > MALE _touch>close_’). However, significant group differences in context-dependent reward variation related to the presence of the female or male experimenter (‘FEMALE > MALE’) revealed that cannabis users displayed altered activity in a cluster encompassing the right dorsal striatum (peak at 27 / 17 / -1, putamen, t^(44)^ = 5.21, k = 87, p = .014) (**Figure 2**). Extracted parameter estimates demonstrated that controls exhibited increased dorsal striatal activity during the presence of the female experimenter relative to the male experimenter (t_(23)_ = 2.71, p = .01, paired t-test), whereas cannabis users exhibited the opposite pattern (t_(21)_ = -4.84, p < .001, paired t-test). The striatal response dynamics mirrored the condition-specific pleasantness experience in the controls, but not in cannabis users (**Figure 2**)

**Figure 2.**
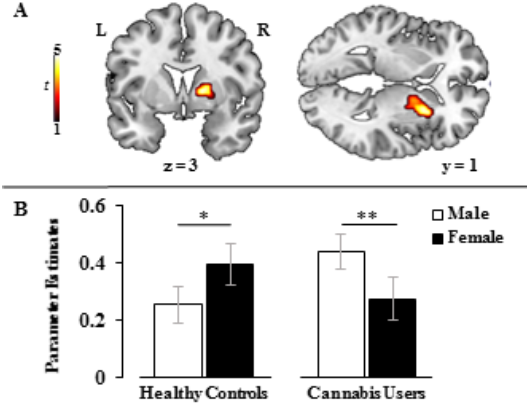
Striatal Response to Rewarding Female Interaction compared between groups. **A:** Difference in striatal activation at MNI-coordinates x = 27 / y = 17 / z = -1 in contrast ‘FEMALE > MALE’ between cannabis users (n = 22) and controls (n = 24) displayed at p_FWE-corrected_ < .05, cluster level. **B:** Extracted parameter estimates from significant cluster from contrasts ‘MALE > Baseline’ (□) and ‘FEMALE > Baseline’ (■) per group. In controls, the striatal response increases significantly upon female interaction. In users, striatal activity decreases. Error bars indicate SEM. *p < .01, **p < .001.

### Associations with Severity of Cannabis Use and Recovery with Abstinence

Measures of withdrawal showed no significant association with behavioral or neural indices (all p > .05). A higher cumulative lifetime use was significantly associated with a stronger decrease in dorsal striatal activity during the presence of the female experimenter relative to the male experimenter (‘FEMALE > MALE’) (r = -.48; p = .024, R^2^ = .23) **(Figure 3),** suggesting an association between a higher cannabis exposure and stronger alterations. The duration of abstinence was not significantly associated with neural indices (p > .24) consistent with the notion that striatal alterations may be enduring rather than transient.

**Figure 3.**
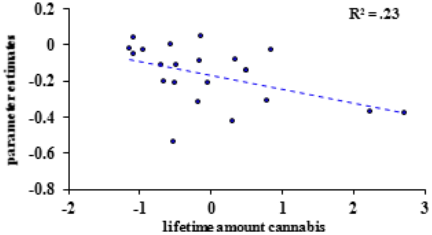
Hedonic Activity and Severity of Cannabis Use. Activation of the dorsal striatum upon ‘FEMALE > MALE’ associates inversely with the cumulative lifetime amount of cannabis use in gram. **(x)** z-transformed cumulative lifetime amount of cannabis use, **(y)** parameter estimates from significant cluster from contrast ‘FEMALE > MALE’, r = -.48, p = .024.

## DISCUSSION

Conceptualizations of drug dependence emphasize the important role of exaggerated striatal responsivity to drug-related rewards and concomitantly blunted sensitivity to natural reinforcers in compulsive drug seeking (Volkow et al., 2012; Koob, 2015). To address whether processing of natural rewards is persistently disrupted in cannabis dependence, the present study examined behavioral and neural responses to social rewards and demonstrated social context-dependent alterations in abstinent cannabis dependent individuals. Specifically, upon female compared to male touch, cannabis users displayed a significantly attenuated increase of reward experience compared to healthy controls. Moreover, while control subjects responded to context-dependent reward variation during female as compared to male presence with an increased dorsal striatal activation, cannabis users displayed the opposite pattern. Examining condition-specific pleasantness ratings and striatal activity revealed a convergent pattern in the controls, whereas the pattern of striatal responses appeared to vary independent of pleasantness experience in users, possibly reflecting blunted striatal coding of reward. Alterations in dorsal striatal reward dynamics increased as a function of cannabis dependence severity. However, neural processing of pleasant touch did not differ between abstinent dependent cannabis users and controls.

The striatum codes both the anticipation and delivery of natural reward (Izuma et al., 2008), including the perception of opposite sex physical attractiveness (e.g. Hahn & Perrett, 2014), and show a high sensitivity to social information (King-Casas et al., 2005). Controls exhibited increasing dorsal striatal activity during the putative presence of the female experimenter and a marked increase in pleasantness experience when they believed the touch was applied by the female relative to the male experimenter. This pattern may reflect either direct natural reward processing associated with the higher perceived attractiveness of the female experimenter or an indirect modulation of the reward response via expectations of opposite sex interaction. Although attractiveness ratings did not differ between the groups, dependent cannabis users demonstrated the opposite dorsal striatal activation pattern and an attenuated increase in pleasantness experience reflecting blunted dynamic coding of context-dependent social reward processing. The findings generally converge with previous reports on residual effects of chronic cannabis use on striatal processing of both, non-drug rewards (Nestor et al., 2010; Jager et al., 2013; Martz et al., 2016) as well as social context information (Gilman et al., 2016) and additionally extend the literature with regard to the following aspects.

First, in line with previous findings (Nestor et al., 2010; Martz et al., 2016), striatal reward processing deficits increased as a function of cannabis exposure indicating these maladaptations may be related to chronic use rather than be a predisposition for cannabis dependence. Furthermore, alterations were observed after prolonged abstinence and therefore may reflect lasting adaptations rather than residual effects of recent cannabis exposure. In the context of accumulating evidence on the relevance of intact striatal reward processing of non-drug rewards (for cannabis dependence see e.g. Yip et al., 2014) and social factors (Nikmanesh et al., 2015) for the long-term success of addiction treatment interventions, the present results appear particularly concerning.

Second, blunted dorsal striatal reward coding was specifically observed during context-dependent reward modulation whereas processing of touch remained intact. These findings argue against general natural reward processing deficits in cannabis users, and rather suggest that striatal processing may be impacted differentially depending on the type of natural reward stimulus, adding to previous reports that alterations across striatal subregions in cannabis users vary with social context (Gilman et al., 2016).

Third, there is ongoing controversy whether chronic cannabis use is associated with lasting striatal neuroadaptations as observed for other drugs of abuse (Curran et al., 2016). Initial findings suggest normal dopamine receptor availability in cannabis users (Urban et al., 2012), whereas more recent studies reported decreased striatal dopamine release capacity (van de Giessen et al., 2017). Moreover, the altered striatal dopaminergic response during early abstinence has been directly linked to anhedonia, and dependence severity (van de Giessen et al., 2017). Therefore, the present findings may be linked to dopaminergic striatal dysfunction, yet also argue for a more complex mechanisms.

Striatal dopaminergic neurotransmission is regulated by the endocannabinoid system (Silveira et al., 2016) and endocannabionoid-mediated adaptations in reward pathways have increasingly been associated with chronic drug dependence (Zlebnik & Cheer, 2016). Animal models suggest a direct association between endocannabinoid transmission in the striatum and hedonic experience of natural, sensory rewards (Mahler et al., 2007). Although homeostatic neuroadaptations in the endocannabinoid system rapidly recover with abstinence (Hirvonen et al., 2012), the present findings may reflect sustained disruptions between subjective hedonic experience and striatal responses, or in the interaction of the endocannabinoid system with other transmitter systems. In the context of previous reports on the contribution of striatal dopamine and endocannabinoid neurotransmission to social reward (Parsons and Hurd, 2015), particularly social play/interaction (Manduca et al., 2016) and expectancy-related modulation of reward (Jubb & Bensing, 2013) the present findings may reflect disruptions in the interplay with the dopaminergic system.

Finally, the ventral striatum has been linked to anticipation of rewards (Schott et al., 2008) while the dorsal striatum encodes reward outcomes (Delgado et al., 2003). Previously, observations regarding reward processing alterations in cannabis users pertained to the ventral portion of the striatum (Nestor et al., 2010; Jager et al., 2013; Martz et al., 2016). However, these studies focused on anticipatory reward phases and non-dependent samples. A shift underlying the control of behavior from the ventral to dorsal part of the striatum has been postulated as a common denominator across substance addictions thought to reflect the transition from voluntary to compulsive behavior (Everitt & Robbins, 2013). As such, the current observation of altered dorsal striatal activation may reflect adaptations in neural mechanisms underlying cannabis dependence.

However, potential limitations should be considered. Abstinence was unsupervised and the cut-off of the immunoassays can only reliably detect cannabis use for a maximum of 15 days (Goodwin et al., 2008). Despite previous literature indicating high reliability of self-reported cannabis use (Martin et al., 1988), we therefore cannot entirely exclude sporadic cannabis use during the abstinence phase as small amounts below the cut-off would solely be detectable in quantitative analyses. To control for effects of tobacco the groups were matched with respect to tobacco use and underwent 1.5h of tobacco abstinence. However, confounding effects related to complex tobacco-cannabis interaction and differences in the time since last use cannot be completely ruled out. Cannabis-withdrawal associated sleep- disturbances may persist for up to 4 weeks, however, sleep disturbances have not been assessed in the present study.

Finally, findings are based on male users. Given the growing evidence for sex-differences in reward-processing in drug using populations future studies are needed to evaluate long-term effects of chronic cannabis use on social reward processing in females.

Taken together, cannabis dependence is associated with lasting adaptions in processing of social rewards. Striatal functioning may be affected differentially across different modalities of reward and future research may need to carefully evaluate different reward dimensions when addressing the striatal system in the context of drug dependence.

## ACKNOWLEDGEMENTS

This work was supported by the National Natural Science Foundation of China (NSFC, 91632117; 31530032), the German Research Foundation (DFG, grant: BE5465/2-1, HU1302/4-1), and an Open Research Fund of the State Key Laboratory of Cognitive Neuroscience and Learning.

## SUPPORTING INFORMATION

### Participants and Study Protocols

### Assessment of Potential Confounders

To control for potential confounding effects of depressive symptom load, attention, attitude towards interpersonal touch and social anxiety all subjects completed the Beck Depression Inventory (BDI-II) (Beck et al., 1996), the d2 test of attention (Brickenkamp and Zillmer, 1998), the social touch questionnaire (STQ) (Wilhelm et al., 2001), the social interaction anxiety scale (SIAS) (Mattick & Clarke, 1998) and the state-trait-anxiety inventory (STAI) (Spielberger, 1989). The positive and negative affect schedule (PANAS) (Crawford & Henry, 2004) was completed before and after the MRI session to control for differences in mood.

### Sample Selection

Following a telephone-based assessment of general study eligibility n = 26 cannabis users and n = 24 male controls were invited for a detailed screening appointment. N = 3 cannabis users were excluded due to too high co-use of other illicit drugs according to the exclusion criteria. 23 abstinent male subjects with cannabis dependence and 24 non-using male controls participated in the study. An initial quality check of the pleasantness ratings revealed that one cannabis user consistently rated male touch with 1 – corresponding to ‘very unpleasant’. A comparably negative perception of male touch was not observed in the other participants (minimum – maximum pleasantness ratings for male touch: cannabis users, 9.55-18.85; controls 8.65-17.6). The unusual negative reaction of this participant to male touch was further confirmed by an outlier analysis (z-value = -3.21, male ratings, within the group of cannabis users). Consequently data from this subject was excluded from all subsequent analyses. This exclusion resulted in a sample size of n = 22 cannabis users and n = 24 control subjects for the final behavioral and fMRI data analysis.

### Group Characteristics

Cannabis users reported greater lifetime experiences with illicit drugs (**Table 1**) than controls. Cannabis, however, was the primary drug of abuse. Cannabis users had abstained from cannabis for a minimum of 28 days; one user reported having used cannabis on one occasion 14 days before the experiment, but was included in the analysis due to his negative urine drug screen on the day of the fMRI examination. Groups were comparable in age, years of education, basal attention, attitude towards interpersonal touch, social anxiety measures, and nicotine and alcohol use (all p > .05, **Table 1**).

### Interpersonal Touch Paradigm Parameters

Standardized tactile stimulation was facilitated through thorough training of the male experimenter prior to the onset of the study, and by signaling the duration of the stimulation to the experimenter via visual cues. The order of the 4 s ‘CLOSE’ and ‘TOUCH’ (20 trials each) conditions was randomized and interleaved with a ‘HOME’ period (4-6 s; mean jitter- time 5 s, 40 trials).

### MRI Data Acquisition

MRI data were acquired on a Siemens 3.0 T TRIO scanner (Siemens, Erlangen, Germany) with a T2*-weighted echo-planar imaging (EPI) sequence (TR = 2500 ms, TE = 30 ms, FoV = 192 mm, flip angle = 90°, voxel size = 2.0 × 2.0 × 3.0 mm^3^, matrix size = 64 × 64, slice thickness = 3.0 mm, 37 axial slices with no gap, 224 whole brain acquisitions oriented along the AC-PC axis for each the male and female session. A high-resolution anatomical reference image was acquired using a T1-weighted mprage sequence (TR = 1660 ms, TE = 2.54 ms, FoV = 256 mm, flip angle = 9°, voxel size = 0.8 × 0.8 × 0.8 mm^3^, matrix size = 256 × 256, slice thickness = 0.8 mm, 208 sagittal slices).

### fMRI Data Preprocessing and Analysis

MRI data was processed and analyzed with Statistical Parametric Mapping12 software (SPM 12, Wellcome Trust Centre for Neuroimaging, London, UK; http://www/fil.ion.ucl.ac.uk/spm) implemented in Matlab (The MathWorks Inc., Natick, MA). The first five volumes of each time-series were discarded to assure T1 equilibration. During realignment, affine transformation was applied to correct for head motion between volumes. In a two-pass procedure images were initially aligned to the first image of the time-series and subsequently realigned to the mean image. Normalization parameters were determined using the T1 image and the segmentation algorithm that combines image registration, tissue classification, and bias correction within the same generative model. Next, normalization parameters were used to spatially normalize the functional time-series to the standard stereotaxic Montreal Neurological Institute (MNI) space template resampled at 3.0 × 3.0 × 3.0 mm. Normalized time-series were smoothed with an 8 mm FWHM Gaussian kernel. The ‘HOME’ condition served as implicit baseline in the first level analysis. To control for movement the head motion parameters were included in the first level matrix.

### Evaluation of the Paradigm and Group-specific Activity in the Touch Network

Examination of the interpersonal touch network in healthy controls on the whole-brain level using the contrast ‘TOUCH > CLOSE’ revealed significant (cluster level FWE-correction, p < .05) activity in the bilateral somatosensory cortex, the bilateral insula, the bilateral dorsal striatum, the left anterior and middle cingulate cortex, and left middle temporal gyrus (**Figure S1, Table S1**) in the control subjects. Marijuana users showed activation in a similar functional network including the bilateral somatosensory cortex, the bilateral insula, the right dorsal striatum, the right anterior cingulate gyrus, the bilateral middle temporal gyrus, and the left precuneus **(Figure S1, Table S1)**.

## SUPPORTING FIGURES and TABLES

**Figure S1.**
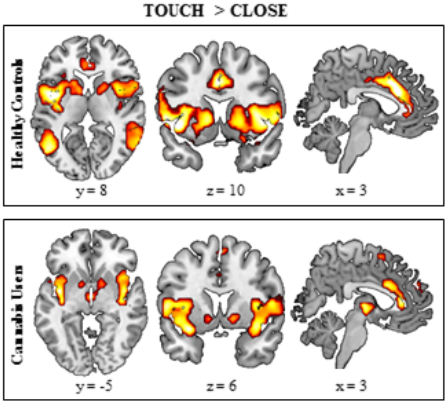
Whole-brain Random Effects Analysis for Contrast ‘TOUCH>CLOSE’ in Controls (n = 24) and Cannabis Users (n = 22). Cluster level FWE-corrected at p < .05, k > 70, MNI-coordinates: x / y / z.

**Table S1.**
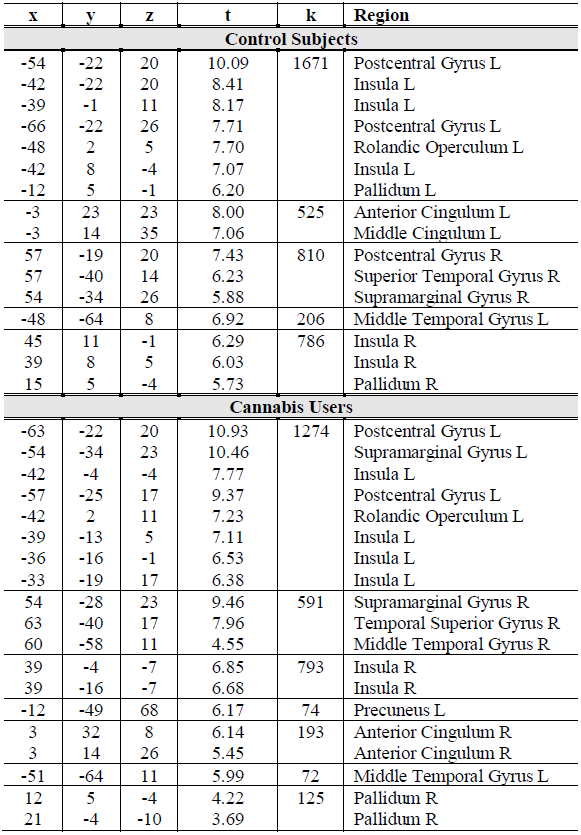
Whole-brain Random Effects Analysis for Contrast ‘TOUCH>CLOSE’ in Controls (n = 24) and Cannabis Users (n = 22). Cluster level FWE-corrected at p < .05, k > 70, MNI-coordinates: x / y / z.

| x | y | z | t | k | Region |
| --- | --- | --- | --- | --- | --- |
| **Control Subjects** |  |  |  |  |  |
| -54 | -22 | 20 | 10.09 | 1671 | Postcentral Gyrus L |
| -42 | -22 | 20 | 8.41 |  | Insula L |
| -39 | -1 | 11 | 8.17 |  | Insula L |
| -66 | -22 | 26 | 7.71 |  | Postcentral Gyrus L |
| -48 | 2 | 5 | 7.70 |  | Rolandic Operculum L |
| -42 | 8 | -4 | 7.07 |  | Insula L |
| -12 | 5 | -1 | 6.20 |  | Pallidum L |
| -3 | 23 | 23 | 8.00 | 525 | Anterior Cingulum L |
| -3 | 14 | 35 | 7.06 |  | Middle Cingulum L |
| 57 | -19 | 20 | 7.43 | 810 | Postcentral Gyrus R |
| 57 | -40 | 14 | 6.23 |  | Superior Temporal Gyrus R |
| 54 | -34 | 26 | 5.88 |  | Supramarginal Gyrus R |
| -48 | -64 | 8 | 6.92 | 206 | Middle Temporal Gyrus L |
| 45 | 11 | -1 | 6.29 | 786 | Insula R |
| 39 | 8 | 5 | 6.03 |  | Insula R |
| 15 | 5 | -4 | 5.73 |  | Pallidum R |
| **Cannabis Users** |  |  |  |  |  |
| -63 | -22 | 20 | 10.93 | 1274 | Postcentral Gyrus L |
| -54 | -34 | 23 | 10.46 |  | Supramarginal Gyrus L |
| -42 | -4 | -4 | 7.77 |  | Insula L |
| -57 | -25 | 17 | 9.37 |  | Postcentral Gyrus L |
| -42 | 2 | 11 | 7.23 |  | Rolandic Operculum L |
| -39 | -13 | 5 | 7.11 |  | Insula L |
| -36 | -16 | -1 | 6.53 |  | Insula L |
| -33 | -19 | 17 | 6.38 |  | Insula L |
| 54 | -28 | 23 | 9.46 | 591 | Supramarginal Gyrus R |
| 63 | -40 | 17 | 7.96 |  | Temporal Superior Gyrus R |
| 60 | -58 | 11 | 4.55 |  | Middle Temporal Gyrus R |
| 39 | -4 | -7 | 6.85 | 793 | Insula R |
| 39 | -16 | -7 | 6.68 |  | Insula R |
| -12 | -49 | 68 | 6.17 | 74 | Precuneus L |
| 3 | 32 | 8 | 6.14 | 193 | Anterior Cingulum R |
| 3 | 14 | 26 | 5.45 |  | Anterior Cingulum R |
| -51 | -64 | 11 | 5.99 | 72 | Middle Temporal Gyrus L |
| 12 | 5 | -4 | 4.22 | 125 | Pallidum R |
| 21 | -4 | -10 | 3.69 |  | Pallidum R |

